# HiG2Vec: Hierarchical Representations of Gene Ontology and Genes in the Poincaré Ball

**DOI:** 10.1101/2020.07.14.195750

**Authors:** Jaesik Kim, Dokyoon Kim, Kyung-Ah Sohn

## Abstract

Knowledge manipulation of gene ontology (GO) and gene ontology annotation (GOA) can be done primarily by using vector representation of GO terms and genes for versatile applications such as deep learning. Previous studies have represented GO terms and genes or gene products to measure their semantic similarity using the Word2Vec-based method, which is an embedding method to represent entities as numeric vectors in Euclidean space. However, this method has the limitation that embedding large graph-structured data in the Euclidean space cannot prevent a loss of information of latent hierarchies, thus precluding the semantics of GO and GOA from being captured optimally. In this paper, we propose hierarchical representations of GO and genes (HiG2Vec) that apply Poincaré embedding specialized in the representation of hierarchy through a two-step procedure: GO embedding and gene embedding. Through experiments, we show that our model represents the hierarchical structure better than other approaches and predicts the interaction of genes or gene products similar to or better than previous studies. The results indicate that HiG2Vec is superior to other methods in capturing the GO and gene semantics and in data utilization as well. It can be robustly applied to manipulate various biological knowledge.

**Availability:** https://github.com/JaesikKim/HiG2Vec

## 1 Introduction

In bioinformatics research, the importance of manipulating biological knowledge has grown, and as a result, how to process knowledge has become more important than just analyzing data alone, for example, KEGG [12, 13, 18, 29], BioCarta [27, 29], Reactome [8, 18, 29], STRING [36, 35, 22, 21], GenBank [2, 26]. Gene Ontology (GO) and Gene Ontology Annotation (GOA), which provide a structured and formal representation of biological knowledge as GO terms, is one of the very large knowledgebases, and there have been several attempts to manipulate and utilize it for analysis. For example, GO enrichment analysis [17, 1, 6, 30], which identifies a common set of GO terms annotated in gene products is widely used in genomic analysis. Another approach for manipulation of GO is to define a semantic similarity measure between GO terms or genes based on relations from GO and GOA, and many ideas have been proposed steadily to date. Information content (IC)-based measures [31, 14, 32, 11, 15, 16, 32, 28, 20] calculate the similarity of two GO terms using the IC of the most informative common ancestor (MICA). GraSM [3] and DiShIn [4] use the average IC of the disjunctive common ancestors (DCAs) instead of the MICA. The Wang method [38] is a graph-based method that calculates the semantic similarity using the topology of the GO graph with hierarchy. Following the introduction of the Wang method, various semantic similarity measures utilizing GO topologies were introduced [41, 10, 19, 23, 7, 42]. Most semantic similarity measures define the semantic similarity between GO terms and further extend it to the semantic similarity between genes through their sets of GO terms from GOA. However, it is difficult to extend these methods into more general applications because only the similarity can be gained.

With the development of deep learning, a concept of word embedding that produces a word representation to understand the semantics of words has been introduced in natural language processing (NLP). It is a mapping of words into vector space to find vector-representation of words using a neural network (NN). Word2Vec [22, 21] is a model for computing better word representations to learn the similarity by using word co-occurrence in a data-driven way. By applying the principle of Word2Vec, Onto2Vec [33] and OPA2Vec [34] were proposed to learn structural information and all relations between GO terms or gene products from GO and GOA. Gene2Vec [5] learned gene co-expression patterns from the Gene Expression Omnibus (GEO) repository. These studies have shown that the Word2Vec-based gene or gene product vector representations can capture gene functions better than conventional semantic similarity measures. From the perspective of knowledge manipulation, vector representation has an advantage of facilitating further analysis based on deep learning [37].

The Word2Vec-based methods used in the previous studies have the advantage of being easy to apply, but they have limitations: when learning the embeddings, they do not consider the latent hierarchy of the large complex graph, which results in the expression of limited GO information [24, 25]. Notably, although the method recognizes the similarity of the GO term, it may be limited in containing the GO hierarchical structure and may not capture hierarchical information, such as the GO term with a higher level containing a more semantically specific concept than with a lower level. In the end, the failure to fully capture GO semantics could affect the embeddings of genes, resulting in incorrect capturing of gene semantics.

In this paper, we propose HiG2Vec, which provides hierarchical vector representation of GO terms and genes by utilizing Poincaré embedding [25] to mitigate the difficulties of hierarchy embedding. Poincaré embedding is a method to learn hierarchy representation on the Poincaré ball, which is a model for hyperbolic geometry in the n-dimensional unit ball. In our method, HiG2Vec proceeds in two steps. First, the model learns representations of GO terms from the GO corpus using Poincaré embedding, and then, it learns representations of genes through fine-tuning using the GOA corpus. We expect that our proposed method captures the GO semantics and gene semantics from GO and GOA better than other methods because of the hierarchical property of GO. We verify the strength of our model by showing several perspectives on GO-level and gene-level evaluations and applying them to various species, namely, humans (*Homo sapiens*), mice (*Mus musculus*), and yeast (*Saccharomyces cerevisiae*). The overview of the proposed HiG2Vec model and our validation schemes are summarized in Figure 1. We share both 200- and 1,000-dimensional vector representations for human, mouse, and yeast. The implementation is available at “https://github.com/JaesikKim/HiG2Vec”.

**Figure 1:**
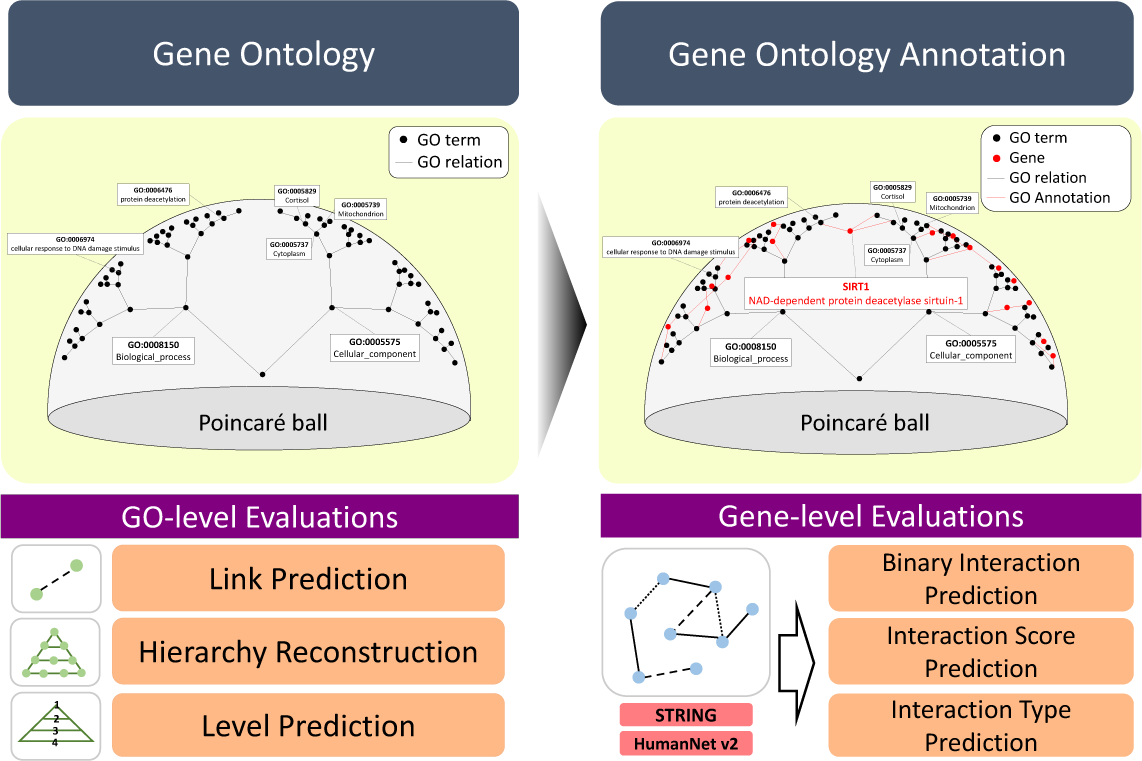
Overview of HiG2Vec. In the n-dimensional Poincaré ball, our model learns vector representations of biological entities first from GO and then from GOA. One example of relations in the 3-dimensional Poincaré ball is shown: SIRT1 and its related GO terms (`GO:0006974`, `GO:0006476`, `GO:0005829`, `GO:0005739`, and `GO:0005737`). As a validation of the results, we performed a total of six experiments at both the GO level and the gene level. We evaluate the representation power of the embedding model by predicting GO links and reconstructing hierarchy and by predicting the binarized interaction, interaction score and interaction type using the STRING and HumanNet v2 databases.

## 2 Related Work

### 2.1 Embedding of GO and Gene/Protein

In addition to methods for obtaining semantic similarities from the partial structure of the ontology, several methodologies for considering the structure of the entire ontology have been introduced with the development of deep learning (Table 1). Inspired by Word2Vec [22, 21] to learn the relations based on the similarity, the embedding model can find a mapping of all biological entities into the vector space. Feature vectors of embedding results can encode both the data structure and the annotations of the entity.

**Table 1:**
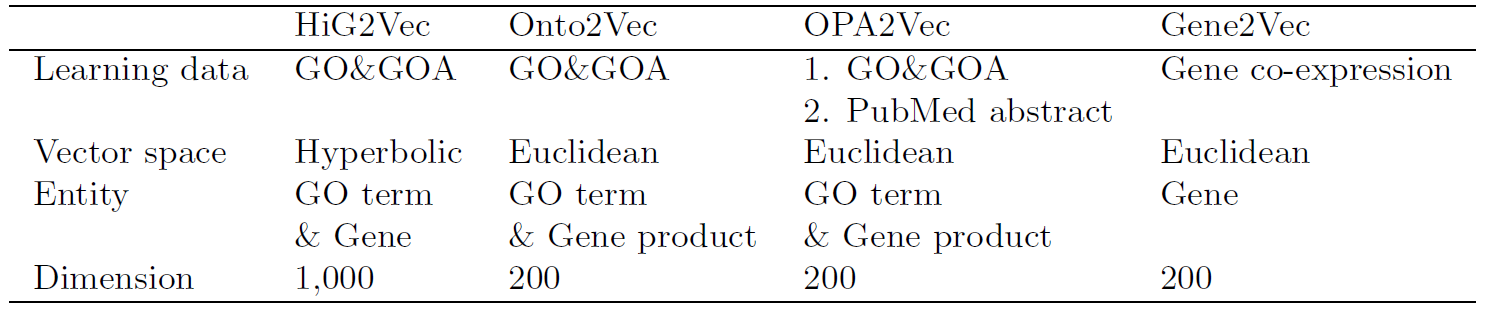
A comparison of the embedding methods

|  | HiG2Vec | Onto2Vec | OPA2Vec | Gene2Vec |
| --- | --- | --- | --- | --- |
| Learning data | GO&GOA | GO&GOA | 1. GO&GOA<br>2. PubMed abstract | Gene co-expression |
| Vector space | Hyperbolic | Euclidean | Euclidean | Euclidean |
| Entity | GO term<br>& Gene | GO term<br>& Gene product | GO term<br>& Gene product | Gene |
| Dimension | 1,000 | 200 | 200 | 200 |

#### 2.1.1 Onto2Vec

To the best of our knowledge, Onto2Vec [33] was the first semantic similarity-based embedding model for gene products as well as GO terms. The corpus that was used for learning representations of entities was generated from GO and GOA. In GO, all types of relationships between GO terms were regarded as *SubClassOf*, and if a GO term was associated with the function of a particular gene in GOA, their relationship was regarded as *AnnotatedWith*. The best result of Onto2Vec was on the 200-dimensional space, and the authors showed that their embedding approaches outperformed traditional semantic similarity measures for relation predictions.

#### 2.1.2 OPA2Vec

Onto2Vec was extended to OPA2Vec [34] by exploiting various aspects of ontologies. The authors incorporated ontology metadata into the Onto2Vec corpus. They considered metadata such as *label, created by, synonym* and *namespace* to be valuable information about classes, relations, and instances. Moreover, they applied pretrained word vectors from Word2Vec on all the PubMed abstracts using transfer learning. They finally showed that their model on the 200-dimensional space had significant improvement compared with Onto2Vec. Both embedding models showed that the function of gene products was captured into their vector representation through various evaluations.

#### 2.1.3 Gene2Vec

Gene2Vec [5] is a method to find the exact functional annotation of genes through the quantitative semantic representation of genes. The authors learned the representation of genes from the gene co-expression patterns as in the Word2Vec-based embedding model. They generated a corpus by extracting human gene expression patterns from the GEO repository. They also showed that their representation in the 200-dimensional space could capture the functional relatedness of genes through predicting gene-gene interactions and recovering known pathways.

## 3 Methods

### 3.1 Poincaré Embedding

Poincaré embedding [25] is a method to embed into an n-dimensional Poincaré ball. This approach finds the optimal embeddings of entities via minimizing the loss value (Supplementary Section 2.3). To embed the entities that contain the hierarchical structure, the transitive closure (Supplementary Section 2.4) of a corpus is used. Poincaré embedding outperforms typical Euclidean embedding in terms of hierarchy representation capacity and generalization ability. The powerful advantage of Poincaré embedding is to successfully preserve hierarchical structure and similarity together.

### 3.2 Learning GO Representations by Poincaré Embedding

Different from the semantic similarity measures or the other embedding methods that considered three domains of GO (BP, MF, and CC) as independent DAGs, we integrated three of them into one DAG using a fake root (Supplementary Section 1.1). The reason is that we wish to embed entire GO terms at the same space as one DAG, not three DAGs. We collected the GO term relations from GO. At this step, we removed the GO term if a GO term was denoted as *is obsolete* and kept the GO term and its alternative GO terms if a GO term was denoted as *alt id*. All the relations of GO became a pair of GO terms in the GO corpus. The GO corpus contained a total of 95,368 relations for 45,026 GO terms. Then, the corpus applied a transitive closure, ultimately resulting in 1,290,646 relations. Let *D*_*GO*_ = {(*GO*_*i*_, *GO*_*j*_)} be a set of relations between GO terms from the GO corpus and {*D*_*GOA*_ = {(*gene*_*i*_, *GO*_*j*_)} be a set of relations between gene and GO terms from the GOA corpus. *B*^*d*^ = {*x* ∈ *R*^*d*^| ‖*x*‖ < 1} is an open *d*-dimensional unit ball, where ‖·‖ denotes the Euclidean norm. Based on equations 1 and 2, we trained our embedding model to find the embeddings 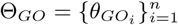, where 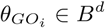.

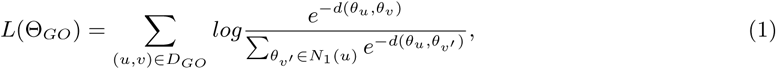

where *N*_1_(*u*) = {*v*|(*u, v*) ∉ *D*_*GO*_} ∪ {*u*} is the set of negative samples for *u*. We randomly sampled 50 negative examples per positive example.

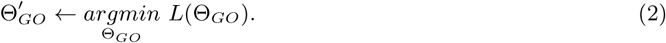

Technically, we used the *burn-in* phase introduced in Poincaré embedding [25], which constitutes small initial epoch training with a reduced learning rate to find better embeddings. The embeddings on 10, 20, 50, 100, 200, 500, and 1,000-dimensional space were trained by the transitive closure of the GO corpus, with a 0.3 learning rate, 50 negatives, 20 burn-in phase, and 1,000 epochs as parameter settings. These embeddings were denoted as HiG2Vec GOonly.

### 3.3 Gene Embedding by Fine-tuning

Subsequently, we organized the GOA corpus for humans, mice, and yeast, consisting of the relations between GO terms and genes. Among annotations in GOA, we removed automatically assigned annotations whose evidence codes were `IEA` or `ND`. As a result, the following evidence codes were selected: `EXP`, `HDA`, `HEP`, `HMP`, `IBA`, `IC`, `IDA`, `IEP`, `IGI`, `IKR`, `IMP`, `IPI`, `ISA`, `ISM`, `ISO`, `ISS`, `NAS`, and `TAS`. Then, we collected pairs of GO terms and genes from the GOA as the GO corpus. If a GO term belonged to the alternate GO terms that we previously kept, we replaced it with its corresponding original GO term. The human GOA corpus ultimately consisted of 17,960 genes and 198,914 relations, the mouse GOA corpus consisted of 19,309 genes and 228,202 relations, and the yeast GOA corpus consisted of 5,997 genes and 40,671 relations. Similarly, we found 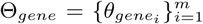 in the same space, where 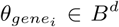, when fine-tuning the embeddings of GO terms together.

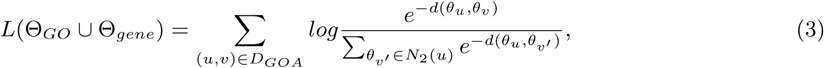

where *N*_2_(*u*) = {*v*|(*u, v*) ∉ *D*_*GOA*_} ∪ {*u*} is the set of negative samples for *u*.

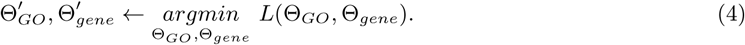

After learning the representation of GO terms sufficiently in the previous step, we fine-tuned the model by the GOA corpus based on pretrained HiG2Vec GOonly with the same parameter settings as before. The genes in GOA were embedded in the same Poincaré ball, while the GO terms were rearranged to be flexible to the relations in both the GO and GOA corpora. These final embeddings were denoted as HiG2Vec. Table 1 summarizes the comparison of HiGVec and other embedding methods. To see the effect of the pretraining, we trained a model that had previously been trained with only GOA information without a pretraining step, denoted as HiG2Vec (non-pretrained).

## 4 Experiments

To verify the strength of HiG2Vec, it was compared to previous embedding methods at the GO level and gene level. By using the GO embeddings, we predicted a link between two GO terms and reconstructed the GO hierarchy, which were previously proposed measures (Nickel2017). In addition, we predicted the level of GO terms in the GO hierarchy using a neural network. By using gene embeddings, we predicted interactions between two genes or gene products from the STRING (Szklarczyk2018) and HumanNet v2 (Hwang2018) databases. Specifically, three kinds of information from interaction were predicted: existence, score, and type.

### 4.1 Link Prediction, Hierarchy Reconstruction, and Level Prediction at the GO Level

To measure the representation power of GO embeddings, we first evaluated generalization performance. The whole GO except the fake root includes 71,639 BP, 13,919 MF, and 7,717 CC links. We sampled as many negative links as positive links: 72,400 BP, 13790 MF, and 7747 CC. Therefore, all samples contained a total of 95,368 positives and 93,937 negatives. If the link was the positive sample, it was labeled as 1; otherwise, it was labeled as 0. Using these link samples, we made GO link predictions [25] of the embedding methods based on the similarity as calculated from the vector representation of GO terms. During this evaluation, the similarity metric used in the evaluation for each model was the same as when training the model. In other words, the embeddings of Onto2Vec and OPA2Vec used cosine similarity, while HiG2Vec GOonly and HiG2Vec used Poincaré similarity (Supplementary Section 2.5).

As another evaluation, the GO hierarchy representation capacity of GO embeddings was measured. We reconstructed all GO relations that were used for learning the embeddings. The reconstruction error was measured for the capacity of the model [25]. Two measures, mRank and mAP, which were obtained as a result of reconstructing the entire transitive closure of relations in GO, can indicate how much the model conserves GO hierarchy. In detail, for each GO term, the other GO terms were ranked based on the distance between them. Then, mRank and mAP were calculated by using the distance-based rank of ancestor GO terms (Supplementary Section 2.6). When calculating the distance, each embedding model used the same distance metric as the metric they used for training. On the graph, the lower the mRank, the closer reachable nodes are, and the larger the mAP, the more accurately reachable nodes are predicted.

Finally, to assess the hierarchical information of GO embeddings, we predicted the level of the GO term through supervised learning. Most GO terms have multiple levels since there is not just one but multiple paths from the root node in the GO DAG structure. To determine one level for every GO term, we chose the maximum level, that is, the longest path among the multiple paths. We designed a simple regression model using a neural network (Supplementary Table S4), whose input is a concatenation of two input vectors and whose output is the predicted level. We tested the model by 10-fold cross-validation. During each fold, we split 25% of the training set as an additional validation set to find a proper learning epoch to help avoid overfitting. For better learning and evaluation, levels with fewer than 10 samples were removed from the dataset. We used the *burn-in* phase and learning rate decay. In addition, the checkpoints of the best model parameters during training were kept, which can be used to choose the best checkpoint for the test set. To measure the prediction results, we used the R-squared and root mean squared error (RMSE) metrics.

### 4.2 Interaction Prediction at the Gene Level

To evaluate the representation power of the models at the gene level, we performed three interaction prediction tasks at the gene level, which compared all the semantic similarity measures and the embedding methods: 1) binary interaction prediction, 2) interaction score prediction and 3) interaction type prediction. Through experiments on different species (human, mouse, and yeast) and various databases (STRING and HumanNet v2) (Supplementary Section 1.2), we evaluated both the robustness of the model and the ability to capture semantics. Among several semantic similarity measures, we selected the Resnik method [31], Wang method [38] and GOGO [42] as representatives of the semantic similarity measure (Supplementary Section 2.2). For the binary and type prediction, we removed relations with confidence scores less than 900 in the STRING database to use relations from interaction databases with the highest confidence [36] for evaluation. In the HumanNet v2 case, because there is no standard threshold for a high LLS, we cut off the relations with scores lower than a certain value (we chose 2 as a threshold). Although the concepts of gene and gene products are not exactly the same, most of the gene functions correspond to the function of the gene product. During our experiments, it is assumed that all genes annotated with GOA can be mapped one-to-one with the proteins in the STRING database in terms of their functions.

#### 4.2.1 Binary Interaction

This task predicts whether there is a relationship between two genes or gene products based on the similarity between them. The thresholded relations were used as positive interactions with labels 1. For the binary interaction prediction, we sampled negative interactions and labeled them as 0. Hence, we obtained four balanced datasets covering three species: 269,785 positives and 269,750 negatives for STRING_Human, 223,140 positives and 224,243 negatives for STRING_Mouse, and 33,069 positives and 33,049 negatives for STRING_Yeast. Similarly, HumanNet_XN consists of 243,083 positives and 247,138 negatives. The interaction between genes or proteins was predicted by their similarity. There were three approaches to calculate the similarity. The best-match average (BMA) computes the similarity between genes from the similarity between GO terms using the best-match average method (Supplementary Section 2.5). The distance-based (DIST) calculates the similarity from the distance between two entities directly. It is applied to the embedding methods only. An NN applies supervised learning to train a similarity measure. It is also applied to embedding methods only. We first split the interaction dataset into 60%, 20%, and 20% for the training, validation, and test sets, respectively. We designed an MLP model (Supplementary Table S4), whose input is a concatenation of two input vectors. The output of the NN can vary from 0 to 1, and this output was used as a predicted similarity. During training, we used the *burn-in* phase and learning rate decay, and the checkpoints to retain the parameter of the best model for the validation set.

#### 4.2.2 Interaction Score

Interaction scores in STRING and HumanNet v2 can be predicted by the NN approach. The dataset was used without removing data by the threshold. Therefore, we gathered 1,159,786 samples for STRING_Human, 1,485,686 samples for STRING_Mouse, 264,653 samples for STRING_Yeast, and 525,515 samples for HumanNet_XN. The same network architecture (Supplementary Table S4) and the same data splitting were used as in the NN approach of binary interaction. After training the prediction model, a coefficient of determination (R-squared) and the mean squared error or RMSE were calculated from the prediction score of the trained network and the ground truth score.

#### 4.2.3 Interaction Type

In STRING, the protein-protein interaction contains one or more types. This interaction type can also be predicted by the NN approach. Considering the type of interaction, we took into account its directionality. For instance, (*A, B*) and (*B, A*) in the STRING database were considered different. After thresholding the data, we ultimately gathered 539,524 samples for STRING_Human, 446,246 samples for STRING_Mouse, and 66,109 samples for STRING_Yeast. This task is a multilabel classification problem, so we used a threshold-dependent neural network (TDNN) [9], which constructs one NN to yield the probabilities for all labels (Supplementary Table S4). We split the dataset into 60%, 20%, and 20% and selected 0.5 for all thresholds of 7 interaction types. The interaction type prediction was performed on the STRING dataset of three species. For measurements of the multilabel classifier, we used the accuracy, macro F1-score, and micro F1-score.

## 5 Result

### 5.1 Link Prediction, Hierarchy Reconstruction, and Level Prediction on the GO Level

In the link prediction experiments, we predicted whether a link exists, given two GO terms, based on the distance between the two GO terms from the entire ontology or each domain. We compared the prediction results of human HiG2Vec and HiG2Vec GOonly with those of human OPA2Vec and human Onto2Vec via receiver operating characteristic (ROC) curves (Figure 2A). Except for the CC domain, the 200-dimensional HiG2Vec performed better than the 1,000-dimensional HiG2Vec. The 200-dimensional HiG2Vec overwhelmingly outperformed Onto2Vec, and the AUC difference from OPA2Vec was only 0.0005 higher for whole domains. A comparison of HiG2Vec and HiG2Vec GOonly indicated that the fine-tuning, during which the positions of GO terms changed as we trained the GOA, degraded the link prediction performance. This occurred because the addition of gene annotation information resulted from the more appropriate adjustment of GO links. In addition, for other species, when comparing the ROC curve of the prediction results of HiG2Vec with that of OPA2Vec or Onto2Vec, the experimental results showed similar trends (Supplementary Figures S1, S3).

**Figure 2:**
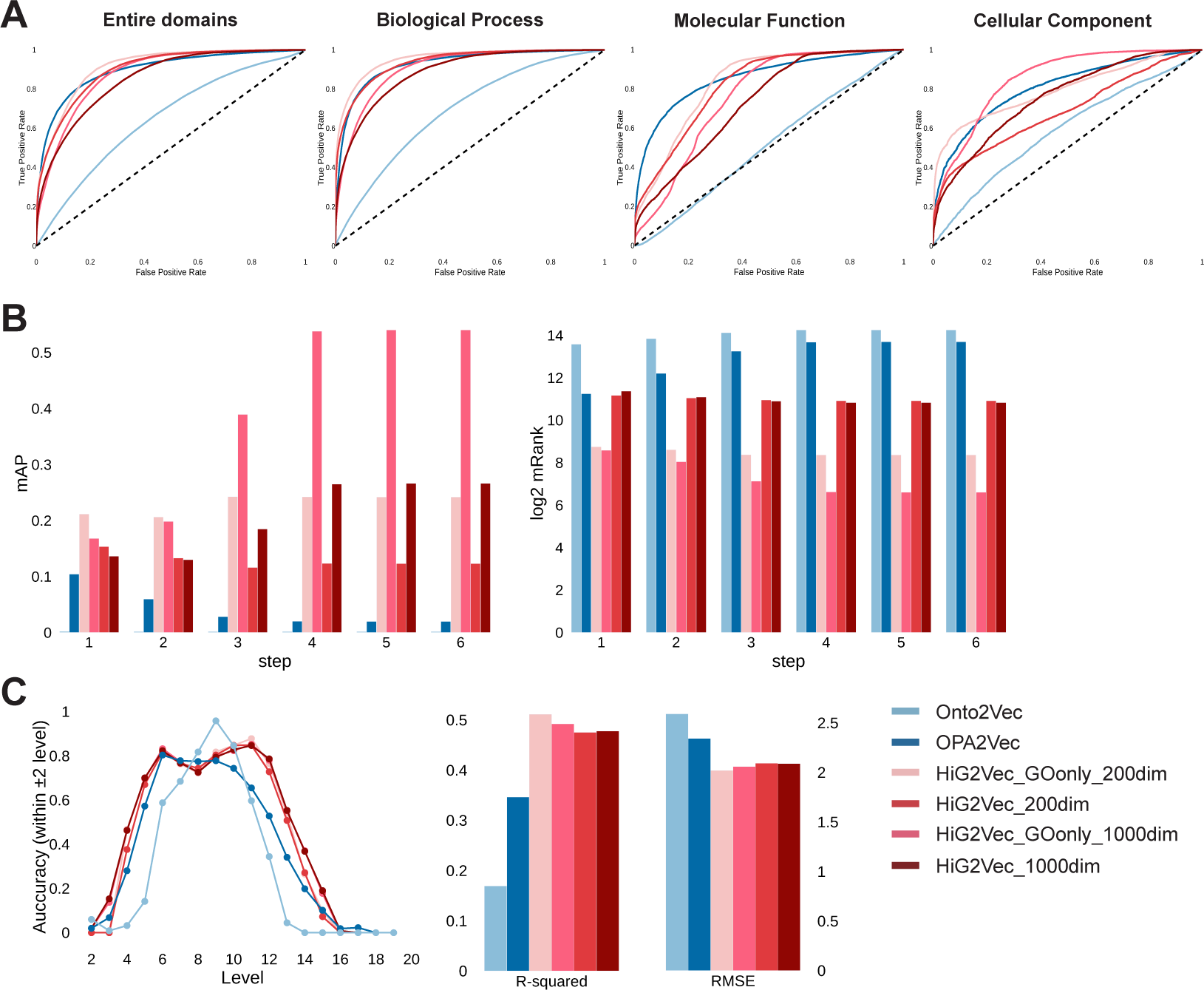
(A) Results of GO link prediction for human GO embeddings: ROC curve using entire domains, using only the BP, MF and CC domains, respectively. (B) Results of GO hierarchy reconstruction for human GO embeddings: log2 transformed mRank and mAP when reconstructing within n-step reachable nodes. (C) Results of GO level prediction for human GO embeddings: prediction accuracy (allowed within 2 level as correct) and their results are evaluated by the R-squared and RMSE metrics.

For the hierarchy reconstruction, we reconstructed not just out-neighbor nodes but also the transitive closure of GO relations step by step so that we could gradually see how the latent hierarchy is preserved in the embeddings. In Figure 2B, the 1,000-dimensional human HiG2Vec and HiG2Vec GOonly are compared with OPA2Vec and Onto2Vec. For reconstructing all *n*-reachable nodes, HiG2Vec GOonly and human HiG2Vec showed a smaller mRank and larger mAP than the others. Fine-tuning our model induced performance degradation for both measures. In particular, reconstructing more reachable nodes resulted in greater performance deterioration, indicating that every node and its distant ancestors underwent a large relocation of positions while reflecting all relations from the GOA. Nevertheless, our models still showed better reconstruction performance than the other embedding models. In addition, experiments with the embeddings of other species showed similar trends. When comparing the mRank and the mAP of mouse HiG2Vec (Supplementary Figure S2) and yeast HiG2Vec (Supplementary Figure S4) with those of OPA2Vec and Onto2Vec, our methods showed better performance for hierarchy reconstruction.

Finally, we predicted the maximum level of GO terms by the human GO embeddings as an input of the NN. For each level, if the difference between the predicted level and the actual level is within two, it is treated as correct in Figure 2C, and the R-squared and RMSE of the prediction results are compared. We can see that the predicted level is closer to the ground truth for HiG2Vec than the other embedding methods for humans. For mouse and yeast, the R-squared and RMSE values of HiG2Vec are better than those of the other embedding methods (Supplementary Table S3 and Supplementary Figure S5). According to the results, our model can better predict the level of GO terms through the information contained in the embedding for the three species.

### 5.2 Interaction Prediction at the Gene Level

#### 5.2.1 Binary Interaction

The similarity between genes for the semantic similarity measures was obtained through BMA (Supplementary Section 2.5). We obtained the similarities between GO terms or genes by the Resnik and Wang method by using the R package GOSemSim [40]. For the embedding models, the similarity was obtained by three approaches: BMA, DIST, and NN. For the total four interaction databases, ROC curves for binary prediction were compared by selecting only the approach with the best performance for each model (Figure 3A).

**Figure 3:**
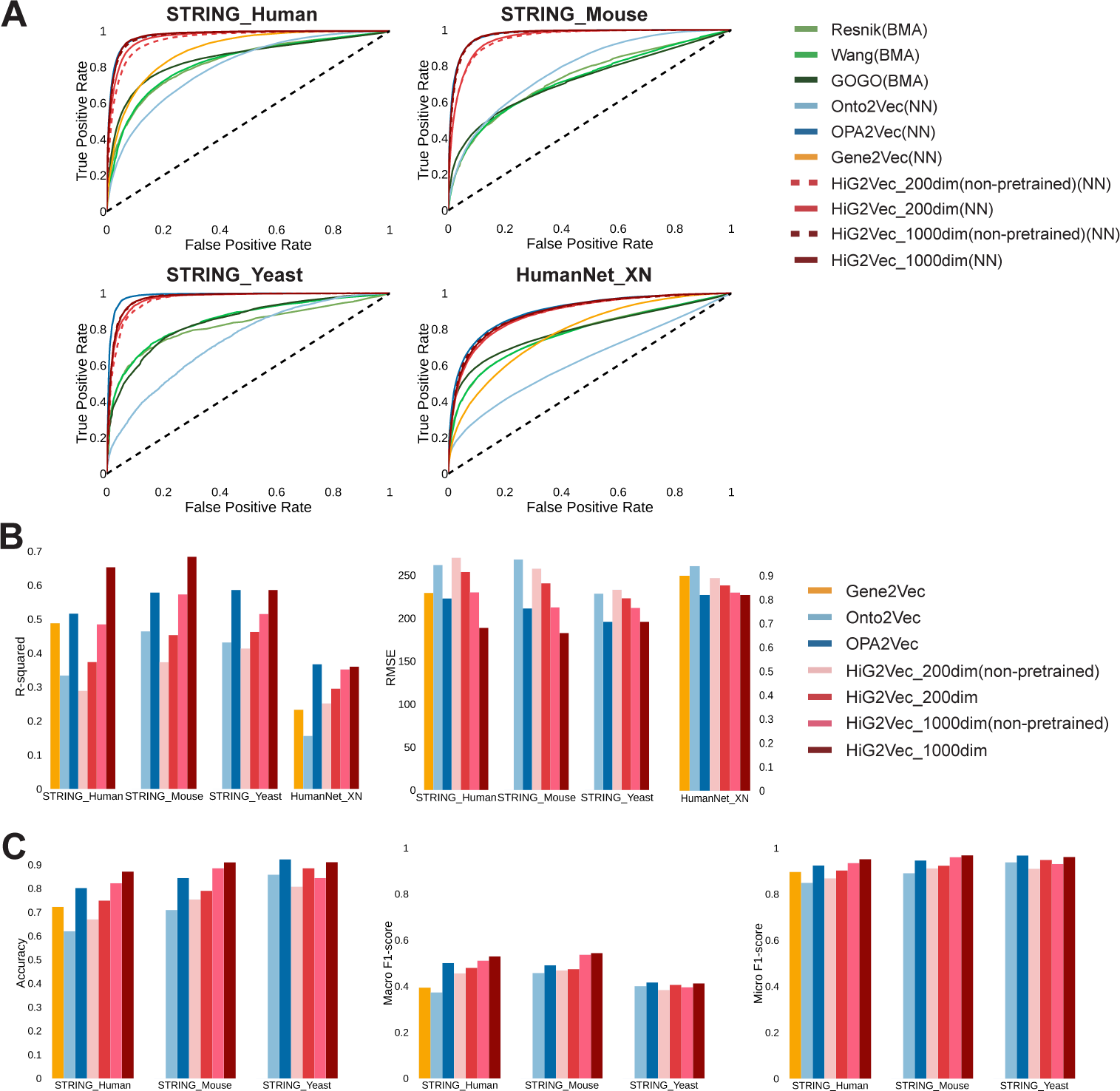
(A) Binary interaction prediction for the species (human, mouse, and yeast) and the databases (STRING and HumanNet v2): ROC curves for humans using STRING_Human, STRING_Mouse and STRING_Yeast, and HumanNet_XN, respectively. (B) Interaction score prediction of the species and the databases: the R-squared and RMSE outputs using STRING_Human, STRING_Mouse, STRING_Yeast, and HumanNet_XN, respectively (C) Interaction type prediction using the STRING database: the accuracy, macro F1-score and micro F1-score, respectively

In the experiments of STRING human protein-protein interaction prediction, Resnik, Wang, and GOGO reached the best performance when using the BP domain compared to using the other GO domains (Supplementary Table S5). In all experimental results of the embedding methods, the NN approach performed overwhelmingly better than either BMA or DIST (Supplementary Table S6). For Gene2Vec, the BMA approach was not available, and the NN approach performed better than the DIST approach. In the case of HiG2Vec, the 1,000-dimensional HiG2Vec could predict interaction better than the 200-dimensional HiG2Vec. Both dimensions of HiG2Vec showed higher performance than each dimensional HiG2Vec (non-pretrained). A comparison of the 1,000-dimensional HiG2Vec with the other embedding methods indicated that the former outperformed Onto2Vec and Gene2Vec but that its performance was slightly lower than that of OPA2Vec (only 0.002 AUC difference). The experiments using STRING_mouse, STRING_yeast, HumanNet_XN showed a similar tendency for the results. For all the embedding methods, the NN approach performed better than the two other approaches (Supplementary Table S6). The 1,000-dimensional HiG2Vec performed better than the 200-dimensional HiG2Vec, and each dimension of HiG2Vec outperformed non-pretrained HiG2Vec. All the cases of HiG2Vec outperformed Onto2Vec, and their performances are minutely lower than that of OPA2Vec, with a 0.0003 AUC difference for STRING_mouse, 0.0126 AUC difference for STRING_yeast, and 0.0156 AUC difference for HumanNet_XN (Supplementary Table S6).

#### 5.2.2 Interaction Score

We predicted the score of the interactions from the STRING and HumanNet v2 databases. The resulting R-squared and RMSE values calculated from the ground truth score and the predicted score are shown in Figure 3B. The performances were obtained from the embedding models in the human, mouse, and yeast STRING database as well as in the HumanNet v2 database. In the results of human and mouse experiments for the STRING database, HiG2Vec showed better performance than HiG2Vec (non-pretrained). The 1,000-dimensional HiG2Vec outperformed all other embedding methods, Gene2Vec (only for humans), Onto2Vec, and even OPA2Vec. However, in the yeast case, OPA2Vec performed slightly better than 1,000-dimensional HiG2Vec, with a difference of 0.0001 for R-squared and 0.04 for the RMSE (Supplementary Table S7). The results of the HumanNet v2 case also showed similar results. Our model outperformed Gene2Vec and Onto2Vec and showed a 0.0073 R-squared difference from OPA2Vec (Supplementary Table S7).

#### 5.2.3 Interaction Type

As a final evaluation, the interaction type provided by STRING was predicted. After training the TDNN-based type prediction model, the test set was evaluated using three metrics: accuracy, macro F1-score and micro F1-score, which were calculated from the ground truth type between the two proteins and their predicted types. The results of comparing the embedding method are shown in Figure 3C. For all species, our model showed better performance on the three metrics than the non-pretrained version and outperformed Gene2Vec (only for humans), Onto2Vec, and OPA2Vec (except yeast). However, the performance difference of STRING yeast experiments between HiG2Vec and OPA2Vec was small: 0.0114 for the accuracy, 0.0038 for the macro-F1 score, and 0.0065 for the micro-F1 score (Supplementary Table S8).

## 6 Discussion

### 6.1 Advantages: Capturing Semantics, Data Utilization, and Robustness

In the above experiments, several methods to capture the semantics of GO terms or genes were compared through various evaluations. The GO embeddings should contain not only GO relations but also information about the hierarchy to better capture the GO semantics. We have shown that HiG2Vec can capture GO semantics through GO-level experiments. When comparing the other embedding methods, our embeddings capture the GO relationship similarly through the GO link prediction, reflect the GO hierarchy better through the hierarchy reconstruction, and contain the hierarchical information better through the level prediction. Next, we showed that understanding the GO semantics better affects capture of the gene semantics by the gene-level experiments. The results of HiG2Vec were superior to those of the semantic similarity measures Onto2Vec and Gene2Vec in terms of capturing the gene semantics by the gene or gene product interaction predictions. In comparison with OPA2Vec, all three gene-level experiments showed comparable performance, in that OPA2Vec uses more knowledge for learning, which ultimately demonstrates the excellence of our model’s methodology. In addition, the experiments of non-pretrained models showed that learning only GO term-gene information without an understanding of GO semantics results in less accuracy in capturing gene semantics. As we hypothesized, HiG2Vec successfully used Poincaré embedding to learn the hierarchy representation of GO, eventually resulting in better capture of the gene semantics.

From the perspective of data utilization, HiG2Vec shows excellent learning efficiency of data information. Our model showed better performance than Onto2Vec despite using the same information and similar overall performance to OPA2Vec despite using less information. In addition, our model performed better than Gene2Vec because GOA already contains enough gene co-expression information, which is the learning data of Gene2Vec. Finally, we conducted learning and STRING protein-protein interaction prediction on each of the other three species. The experiment was also carried out using another interaction database, namely, HumanNet v2. The rankings of all methods varied slightly across species and databases, but the experimental results tended to be consistent. This implies that HiG2Vec can be applied to other species or knowledge bases based on deep learning approaches.

### 6.2 Dimensionality

The decision of dimensionality is difficult but crucial for future applications. The problem of dimensional decision involves a tradeoff. According to the word embedding that has been studied, low-dimensional space is not expressive enough to capture the entire relation, and high-dimensional space has powerful representation ability but is susceptible to overfitting [39]. In addition, because of the large number of parameters, high dimensionality causes an increase in model complexity, which limits model applicability. In this study, performance changes for each experiment were analyzed according to the dimensionality changes. In the link prediction, we noted that the performance decreased as the dimensionality increased (Supplementary Table S1). The hierarchy reconstruction and level prediction experiment showed better performance as the dimension increased (Supplementary Table S2,S3). In the interaction prediction experiments, the performance improved as the dimension increased (Supplementary Table S6-S8). Except for GO link experiments, it seems to be favorable to select higher dimensions in all experiments. Considering the computational cost together, we concluded that 1,000 dimensions are appropriate for overall aspects. We share both 200-dimensional HiG2Vec and 1,000-dimensional HiG2Vec for future work.

### 6.3 Two-step Procedure

Our proposed idea proceeded in two steps of learning: learning GO and learning GOA. The reason is, first, not to limit genes to only the leaf nodes of the hierarchy. If we trained GO and GOA using Poincaré embedding at one time, there would be a constraint on the representation of genes that is restricted to leaf nodes only. Second, there is a computational limitation to learning GO and GOA at once because there are too many relations in the transitive closure of the GOA relation. Therefore, we used only the transitive closure of GO relations for learning GO and did not use the transitive closure for learning GOA. As a result, we expect that the hierarchy was slightly rearranged, making the gene representations more accurate. According to the GO-level evaluation results, the link prediction results showed that our model performed slightly worse after fine-tuning. The results of hierarchy reconstruction also showed that the performance difference between HiG2Vec GOonly and HiG2Vec widened as the number of reachable node steps increased. It can be said that the structure of the hierarchy was disrupted by fine-tuning, as we expected. The GO level prediction results also showed that, as expected, the loss of hierarchy resulted in a slight decrease in the prediction performance compared to that of HiG2Vec GOonly.

## 7 Conclusion

In this paper, we proposed a novel embedding model from GO and GOA by adopting representation learning that specializes in the GO hierarchy. We confirmed that the hierarchical structure was well represented in our model through the GO-level evaluations, and our model showed better representation power than the existing semantic similarity measures and the embedding methods through the gene-level evaluations. The vector representation of GO terms and genes in HiG2Vec is expected to be used for various downstream analyses such as data processing at the gene and protein levels or manipulation of biological knowledge. The fact that the embedding of our model is on the Poincaré ball can limit its application in Euclidean space, but it can be applied to general machine learning or deep learning based applications. To further improve the embedding result, the training corpus can be extended to other domains or other ontologies. In future work, we plan to apply our model to integrate and learn multi-ontology.

