## Supplementary materials for "HiG2Vec: Hierarchical Representations of Gene Ontology and Genes in the Poincaré Ball"

### Supplementary material

#### 1 Materials

##### 1.1 Gene Ontology (GO) and Gene Ontology Annotation(GOA)

The Gene Ontology Consortium [4] provides a structured and formal representation of biological knowledge, describing gene and gene product attributes across all eukaryotes, as described by GO and GOA. The consortium defines a dynamic, controlled vocabulary of all biological knowledge to explain gene and protein roles in all eukaryotes. GO terms are classified in three domains: biological process (BP), molecular function (MF), and cellular component (CC). Each GO term has four types of relations to others: *is a*, *part of*, *has part*, or *regulates* (including *positively regulates* and *negatively regulates*). Each GO domain can be represented in DAG format, where nodes represent GO terms and directed edges represent the relation between GO terms. To integrate three domains into one DAG for further steps, we made a fake root (`GO:0000000`) to connect to the root nodes of three domains. The GOA Database, which is also provided by the Gene Ontology Consortium, provides the knowledge of genes in specific species via GO terms. All annotations include an evidence code to support the relationship between GO term and gene, according to how annotation was gained, such as by an experiment, phylogenetic inference, computational analysis, author statement, electronic annotation, or curator statement.

##### 1.2 Interaction Database

STRING [17] is a database of known and predicted protein-protein interactions. This database has collected interaction information from more than 2,000 species from five main sources: genomic context predictions, high-throughput lab experiments, co-expression, automated textmining, and previous knowledge in databases. In the database, interaction types are classified into 7 types: activation, binding, catalysis, expression, inhibition, post-translational modification, and reaction. The interaction also has a confidence score, which indicates the approximate probability that a predicted link exists (ranging from 0 to 1,000). We chose three species to evaluate the interaction predictors: human (*H. sapiens*), mouse (*M. musculus*) and yeast (*S. cerevisiae*).

HumanNet v2 [5] provides a human gene network for disease research, integrating diverse types of data information, such as protein-protein interaction, co-citation, co-essentiality, co-expression, pathway database, protein domain profile associations, gene neighborhood, phylogenetic profile association, and interlogs from other species, including 17,929 genes and 525,537 links. A link between genes has negative log-likelihood scores (LLSs) that measure the probability of interaction. They provide various versions of the gene-gene network. Among them, we chose to use HumanNet_XN (fully extended network) because it is recommended for studies requiring the most comprehensive networks.

#### 2 Methods

##### 2.1 GO version and processing

Gene Ontology can be downloaded from 'http://geneontology.org/docs/download-ontology/'. For our study, we used go.obo (version 1.2, released/2019-02-27). These files contain the core GO ontology not filtered relationship, including *has_part*. Therefore, the processed GO corpus contains a small number of cycles in the graph, which makes it impossible to obtain the GO level, so we removed one of part_of or has_part link randomly in these cases.

#### 2.2 Semantic similarity measures

##### 2.2.1 Resnik method [15]

The information content of a GO term is calculated by the negative log probability of its occurrence, which is defined as the frequency of the term in GO corpus. Resnik method is defined as:

$$IC(t) = -log(freq(t)) \tag{1}$$

$$sim_{Resnik}(t_1, t_2) = IC(MICA), \tag{2}$$

where the most informative common ancestor (MICA) of two GO terms denotes their closest common ancestor term.

##### 2.2.2 Wang method [18]

Given a GO term $A$, the DAG of $A$ and its ancestors are defined as $DAG_A = (A, T_A, E_A)$, where $T_A$ denotes the set of GO terms in $DAG_A$, and $E_A$ denotes the set of edges in $DAG_A$. The S-value of a GO term in $DAG_A$, which infers the contribution of a GO term $t$ to the semantics of GO term $A$, is defined as:

$$S_A(t) = \begin{cases} 1 & \text{if } t = A \\ max\{w_e * S_A(t')|t' \in children(t)\} & \text{if } t \neq A, \end{cases} \tag{3}$$

where $w_e$ is the semantic contribution factor for edge $e \in E_A$ $(0 < w_e < 1)$. Given $DAG_A$ and $DAG_B$ for GO terms $A$ and $B$ respectively, the semantic similarity between two terms is defined as:

$$sim_{Wang}(A, B) = \frac{\sum_{t \in T_A \cap T_B} (S_A(t) + S_B(t))}{\sum_{t_A \in T_A} S_A(t_A) + \sum_{t_B \in T_B} S_B(t_B)} \tag{4}$$

##### 2.2.3 GOGO [20]

The similarity of GO terms in GOGO is calculated in the similar way as the Wang method. The additional operation in the methodology is to assign a semantic contribution factor ($w_e$). In GOGO, the semantic contribution factor is defined as:

$$w_e = 1/(c + nc(t)) + d, \tag{5}$$

where $nc(t)$ is the total number of children for GO term $t$, and $c$ and $d$ are constant parameter. In practice, $c$ is assigned 0.67, and $d$ is assigned as 0.4, 0.3, and 0.2 for *is a*, *part of*, and *regulates*, respectively.

#### 2.3 Poincaré Embedding

A Poincaré ball is a model of hyperbolic space with constant negative curvature. Hyperbolic space is a non-Euclidean space with negative curvature, where the amount of deviation from planarity is negative. The Poincaré ball is well suited for modeling hierarchies due to the geometric property, in which the distance of points in hyperbolic space increases exponentially the closer they are to the boundary of the unit ball because of negative curvature [7, 13]. In the open $d$-dimensional unit ball $B^d = \{x \in R^d | \, \|x\| < 1\}$, where $\|\cdot\|$ denotes the Euclidean norm, the distance between points $\boldsymbol{u}, \boldsymbol{v} \in B^d$ is given as:

$$d(\boldsymbol{u}, \boldsymbol{v}) = arcosh\left(1 + 2\frac{\|\boldsymbol{u} - \boldsymbol{v}\|^2}{(1 - \|\boldsymbol{u}\|^2)(1 - \|\boldsymbol{v}\|^2)}\right) \tag{6}$$

Equation 6 allows the root node of the tree to lie at the origin $B^d$ and the leaf nodes to lie close to the boundary of the Poincaré ball. Therefore, this framework can capture the hierarchy of objects through their norm and their similarity through their distance.

Poincaré embedding [13] is a method to embed into an n-dimensional Poincaré ball. This approach finds the optimal embeddings of entities by minimizing a distance-based loss value via the following function in Equation 7. $D = \{(u, v)\}$ is a set of relations of objects, and the loss function is defined as:

$$L(\Theta) = \sum_{(u,v) \in D} log \frac{e^{-d(\boldsymbol{u}, \boldsymbol{v})}}{\sum_{\boldsymbol{v'} \in N(u)} e^{-d(\boldsymbol{u}, \boldsymbol{v'})}}, \tag{7}$$

where $N(u) = \{v|(u,v) \notin D\} \cup \{u\}$ is the set of negative samples for $u$. To find embeddings $\Theta = \{\theta_i\}_{i=1}^n$, where $\theta_i \in B^d$, they solve the optimization problem (Equation 8) via the stochastic Riemannian optimization method.

$$\Theta' \leftarrow \underset{\Theta}{argmin}\ L(\Theta) \qquad s.t.\ \forall \theta_i \in \Theta : \|\theta_i\| < 1 \tag{8}$$

Using Riemannian stochastic gradient descent (RSGD) [1], the parameters are updated following Equation 9. Let $\nabla_R$ denote the Riemannian gradient of $L(\theta)$, and let $\nabla_E$ denote the Euclidean gradient of $L(\theta)$.

$$\theta_{t+1} \leftarrow \theta_t - \eta_t \nabla_R L(\theta_t), \tag{9}$$

where $\eta_t$ denotes the learning rate at time t. To derive the Riemannian gradient from the Euclidean gradient, $\nabla_E$ is rescaled by using the inverse of the Poincaré ball metric tensor ($(2/1 - \|x\|^2)^2 g^E$, where $g^E$ denotes the Euclidean metric tensor) since the direction of gradient is identical. The final update rule is as follows:

$$\text{proj}(\theta) = \begin{cases} \theta / \|\theta\| - \epsilon & if\ \ \|\theta\| \geq 1 \\ \theta & otherwise, \end{cases} \tag{10}$$

where $\epsilon$ is a small constant.

$$\theta_{t+1} \leftarrow \text{proj}\left(\theta_t - \eta_t \frac{(1 - \|\theta_t\|^2)^2)}{4} \nabla_E\right) \tag{11}$$

#### 2.4 Transitive Closure

In graph theory, transitive closure of a graph is a graph which contains an $edge(vertice_i, vertice_j)$ whenever there is a directed path from $vertice_i$ to $vertice_j$ [16]. Therefore, the transitive closure of a directed graph indicates its reachability relation. We will use a concept of *n-step reachable*, which means that there is a directed path from $vertice_i$ to $vertice_j$ within $n$ edges. In other words, 1-step reachable node means out-neighbors, while the whole step reachable node means the out-neighbors in transitive closure.

#### 2.5 Similarity Measurement

- **Best-Match Average** The semantic similarities between genes can be computed from the idea of mixing the similarity of GO term pairs, such as Average (Avg) [8], Maximum (Max) [6], Average Best-Matches (ABM) [14, 10] and Best-match Average (BMA) [18, 6]. Among them, BMA was reported as the best approach [20]. The formulas of BMA and ABM were used interchangeably in previous studies [18, 14, 6, 2, 10, 11, 12, 3, 20]. We therefore followed the definition in GOSemSim [19]. Given two genes $gene_1$ and $gene_2$ annotated with $\{GO_{11}, GO_{12}, ..., GO_{1m}\}$ and $\{GO_{21}, GO_{22}, ..., GO_{2n}\}$ respectively, the BMA similarity defined as:

$$BMA(gene_1, gene_2) = \frac{\sum_{i=1}^m \underset{1 \leq j \leq n}{max}\ sim(GO_{1i}, GO_{2j}) + \sum_{j=1}^m \underset{1 \leq i \leq m}{max}\ sim(GO_{1i}, GO_{2j})}{m+n} \tag{12}$$

- **Cosine similarity** In the embedding methods based on Word2Vec in the Euclidean space, cosine similarity are used for calculating similarity between vectors.

$$cosine\_similarity(\boldsymbol{u}, \boldsymbol{v}) = \frac{\boldsymbol{u} \cdot \boldsymbol{v}}{\|\boldsymbol{u}\|\ \|\boldsymbol{v}\|} \tag{13}$$

- **Poincaré similarity** There is no concept of similarity in hyperbolic space [9]. Instead, we applied a monotonic decreasing function to poincaré distance to invert the rank order of distance, as it would play a role of similarity. The following function was used to our embeddings at the evaluation steps that required the similarity of vectors.

$$poincaré\_similarity(\boldsymbol{u}, \boldsymbol{v}) = \frac{1}{1 + d(\boldsymbol{u}, \boldsymbol{v})}, \tag{14}$$

where $d(\boldsymbol{u}, \boldsymbol{v})$ is a poincaré distance between $\boldsymbol{u}$ and $\boldsymbol{v}$.

#### 2.6 Evaluation Metrics

- **AUC**, a Receiver operating characteristic (ROC) curve is widely used for evaluating prediction models. It plots True Positive Rate (TPR) against False Positive Rate (FPR).

$$TPR = \frac{TP}{TP + FN} \tag{15}$$

$$FPR = \frac{FP}{FP + TN}, \tag{16}$$

where TP, FP, TN and FN are the number of true positives, false positives, true negatives, and false negatives respectively. The AUC of ROC stands for the area under the ROC curve. This evaluation metric allow us to compare the prediction models more formally and precisely.

- **mRank**, a mean rank (mRank) of DAG is defined as:

$$N = \sum_{i}^{n} \texttt{count}(nbr(i)) \tag{17}$$

$$mRank = \frac{1}{N} \sum_{i} \sum_{j \in nbr(i)} (obs\_rank_i(j) - exp\_rank_i(j)), \tag{18}$$

where $nbr(i)$ denotes the out-neighbors of vertice $i$, and $obs\_rank_i(j)$ and $exp\_rank_i(j)$ denote an observed and expected distance-based rank of vertice $j$ respectively from vertice $i$.

- **mAP**, a mean Average Precision (mAP) of DAG is defined as:

$$mAP = \frac{1}{n} \sum_{i} AP\_score(i) \tag{19}$$

where $AP\_score(i)$ denotes an average precision score of the rankings from vertice $i$.

### 3 Results

Table S1: Results of GO link prediction for GO embeddings with dimensionality changes

| | | Dim | GOonly BP | GOonly MF | GOonly CC | GOonly All | Human BP | Human MF | Human CC | Human All |
|---|---|---|---|---|---|---|---|---|---|---|
| AUC | HiG2Vec | 10 | **0.9631** | **0.9317** | 0.8320 | **0.9310** | 0.9473 | **0.8965** | 0.6810 | 0.9153 |
| | | 20 | 0.9623 | 0.9134 | 0.8288 | 0.9284 | **0.9489** | 0.8947 | 0.7235 | **0.9167** |
| | | 50 | 0.9590 | 0.8869 | 0.8179 | 0.9219 | 0.9436 | 0.8656 | 0.7120 | 0.9089 |
| | | 100 | 0.9592 | 0.8639 | 0.7859 | 0.9184 | 0.9447 | 0.8463 | 0.6573 | 0.9068 |
| | | 200 | 0.9532 | 0.8332 | 0.7973 | 0.9114 | 0.9362 | 0.8188 | 0.6875 | 0.8979 |
| | | 500 | 0.9243 | 0.7843 | **0.8616** | 0.8926 | 0.9032 | 0.7553 | 0.7584 | 0.8688 |
| | | 1,000 | 0.9065 | 0.7628 | 0.8516 | 0.8779 | 0.8854 | 0.7299 | 0.7568 | 0.8519 |
| | OPA2Vec | 200 | - | - | - | - | 0.9268 | 0.8566 | **0.8064** | 0.8974 |
| | Onto2Vec | 200 | - | - | - | - | 0.6878 | 0.5076 | 0.5886 | 0.6482 |

| | | Dim | Mouse BP | Mouse MF | Mouse CC | Mouse All | Yeast BP | Yeast MF | Yeast CC | Yeast All |
|---|---|---|---|---|---|---|---|---|---|---|
| AUC | HiG2Vec | 10 | **0.9437** | **0.9085** | 0.7058 | **0.9144** | **0.9598** | **0.9218** | 0.7699 | **0.9276** |
| | | 20 | 0.9429 | 0.8898 | 0.7085 | 0.9120 | 0.9590 | 0.9033 | 0.7700 | 0.9252 |
| | | 50 | 0.9009 | 0.7858 | 0.6450 | 0.8615 | 0.9551 | 0.8758 | 0.7318 | 0.9194 |
| | | 100 | 0.9402 | 0.8380 | 0.6509 | 0.9019 | 0.9554 | 0.8515 | 0.6954 | 0.9156 |
| | | 200 | 0.9317 | 0.8168 | 0.6877 | 0.8939 | 0.9491 | 0.8293 | 0.7333 | 0.9086 |
| | | 500 | 0.9039 | 0.7602 | 0.7582 | 0.8693 | 0.9180 | 0.7742 | **0.8061** | 0.8844 |
| | | 1,000 | 0.8777 | 0.7216 | 0.7548 | 0.8446 | 0.8977 | 0.7459 | 0.8011 | 0.8656 |
| | OPA2Vec | 200 | 0.9330 | 0.8680 | **0.8122** | 0.9067 | 0.9129 | 0.8494 | 0.7936 | 0.8793 |
| | Onto2Vec | 200 | 0.6915 | 0.5053 | 0.5879 | 0.6502 | 0.6744 | 0.4895 | 0.5737 | 0.6317 |

Table S2: Results of hierarchy reconstruction for GO embeddings with dimensionality changes

| | Dim | GOonly mRank | GOonly mAP | Human mRank | Human mAP | Mouse mRank | Mouse mAP | Yeast mRank | Yeast mAP |
|---|---|---|---|---|---|---|---|---|---|
| HiG2Vec | 10 | 426.23 | 0.1573 | 2382.90 | 0.0879 | 2160.50 | 0.0718 | 998.93 | 0.1003 |
| | 20 | 364.60 | 0.2129 | 2461.74 | 0.0851 | 2028.34 | 0.0874 | 917.90 | 0.1166 |
| | 50 | 345.29 | 0.2321 | 2250.94 | 0.0952 | 4580.02 | 0.1309 | 748.54 | 0.1695 |
| | 100 | 781.70 | 0.1736 | 2692.95 | 0.0873 | 2700.92 | 0.0880 | 1323.09 | 0.1230 |
| | 200 | 330.30 | 0.2418 | 1928.51 | 0.1228 | 1960.86 | 0.1155 | 772.65 | 0.1667 |
| | 500 | 120.47 | 0.4434 | 1695.99 | 0.2037 | 1926.90 | 0.2083 | 513.94 | 0.3231 |
| | 1,000 | **97.43** | **0.5492** | **1813.38** | **0.2661** | **1911.06** | **0.2422** | **510.35** | **0.3914** |
| OPA2Vec | 200 | - | - | 13257.09 | 0.0197 | 13268.92 | 0.0203 | 10404.58 | 0.0177 |
| Onto2Vec | 200 | - | - | 19531.94 | 0.0014 | 20979.47 | 0.0014 | 16655.53 | 0.0016 |

Table S3: Results of GO level prediction for GO embeddings with dimensionality changes

| | Dim | GOonly R-squared | GOonly RMSE | Human R-squared | Human RMSE | Mouse R-squared | Mouse RMSE | Yeast R-squared | Yeast RMSE |
|---|---|---|---|---|---|---|---|---|---|
| HiG2Vec | 10 | 0.2539 | 2.4924 | 0.2178 | 2.5539 | 0.2341 | 2.5271 | 0.2368 | 2.5226 |
| | 20 | 0.3217 | 2.3781 | 0.3361 | 2.3528 | 0.3291 | 2.3652 | 0.3317 | 2.3606 |
| | 50 | 0.4503 | 2.1409 | 0.4378 | 2.1652 | 0.4075 | 2.2228 | 0.4395 | 2.1619 |
| | 100 | 0.5078 | 2.0258 | 0.4645 | 2.1132 | 0.4534 | 2.1349 | 0.4824 | 2.0776 |
| | 200 | **0.5113** | **2.0187** | 0.4751 | 2.0922 | 0.4635 | 2.1150 | **0.4950** | **2.0519** |
| | 500 | 0.4958 | 2.0504 | 0.4664 | 2.1093 | **0.4866** | **2.0691** | 0.4810 | 2.0802 |
| | 1000 | 0.4922 | 2.0577 | **0.4776** | **2.0871** | 0.4820 | 2.0784 | 0.4872 | 2.0678 |
| OPA2Vec | 200 | - | - | 0.3463 | 2.3421 | 0.3526 | 2.3352 | 0.3172 | 2.3697 |
| Onto2Vec | 200 | - | - | 0.1960 | 2.5902 | 0.2025 | 2.5798 | 0.1820 | 2.6127 |

Table S4: A detail of neural network architectures of interaction prediction.

| Layer type | | Size of output | Remarks |
|---|---|---|---|
| | Input | $2 * d$ | Concatenation of two embedding vectors |
| Layer 1 | Fully Connected | $d$ | $2 * d \to d$ |
| | BatchNorm | $d$ | |
| | ReLU | $d$ | |
| Layer 2 | Fully Connected | GO level prediction: 1 | $d \to 1$ |
| | | Binary prediction: 1 | $d \to 1$ |
| | | Score prediction: 1 | $d \to 1$ |
| | | Type prediction: `count(types)` | $d \to$ `count(types)` |

* $d$ denotes a dimensionality of embeddings

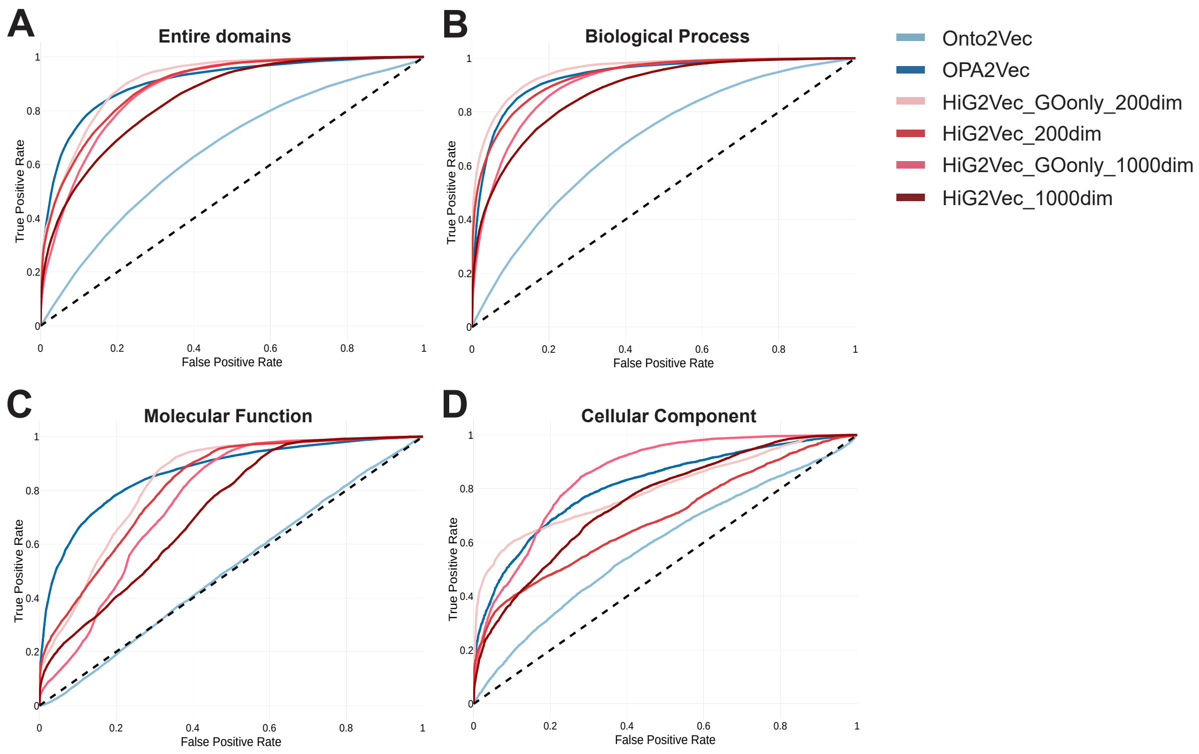


Figure S1: GO Link prediction for embeddings of mouse. (A) is a ROC curve using entire domains and each (B),(C) and (D) are ROC curves when using only biological process (BP), molecular function (MF) and cellular component (CC) domain respectively.

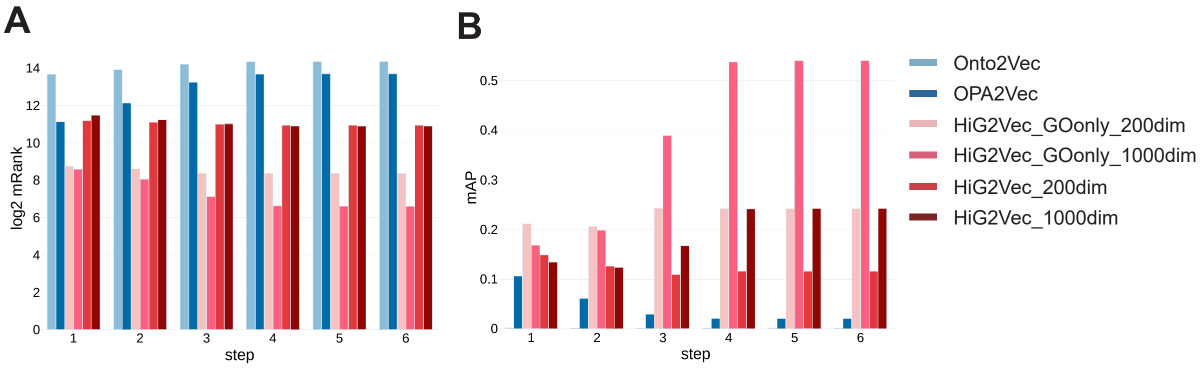


Figure S2: Hierarchy reconstruction for embeddings of mouse. (A) is log2 transformed mRank and (B) is mAP when reconstructing within n-step reachable nodes.

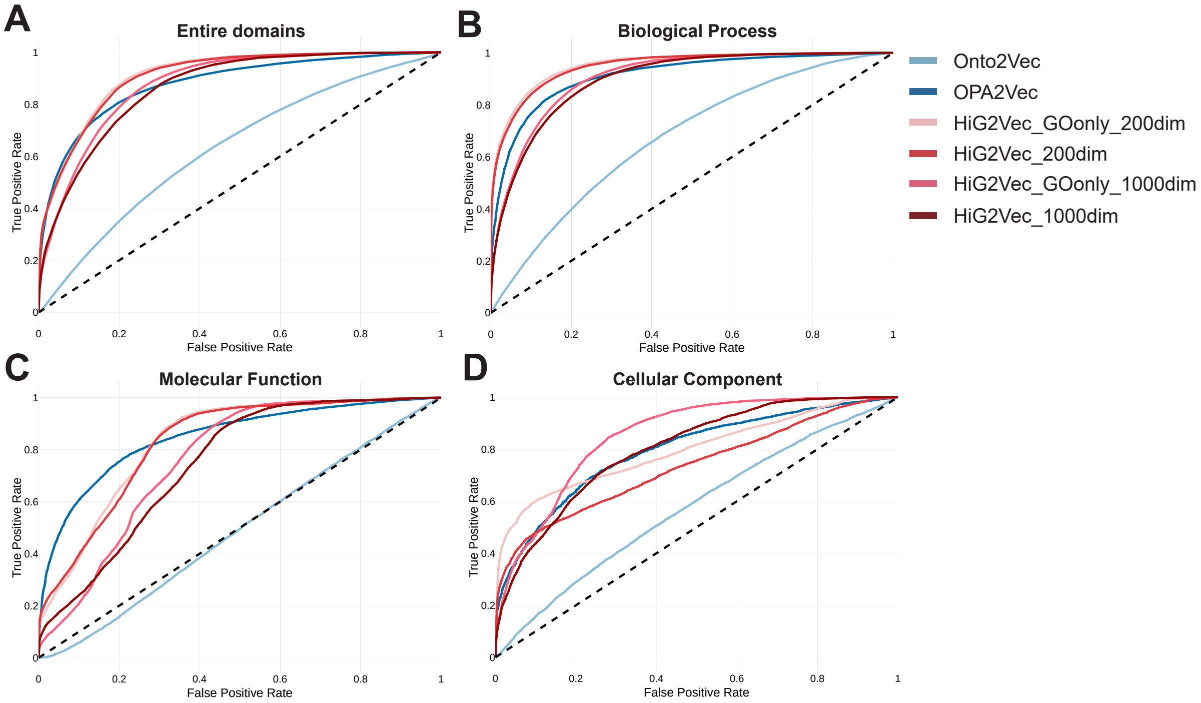


Figure S3: GO Link prediction for embeddings of yeast. (A) is a ROC curve using entire domains and each (B),(C) and (D) are ROC curves when using only biological process (BP), molecular function (MF) and cellular component (CC) domain respectively.

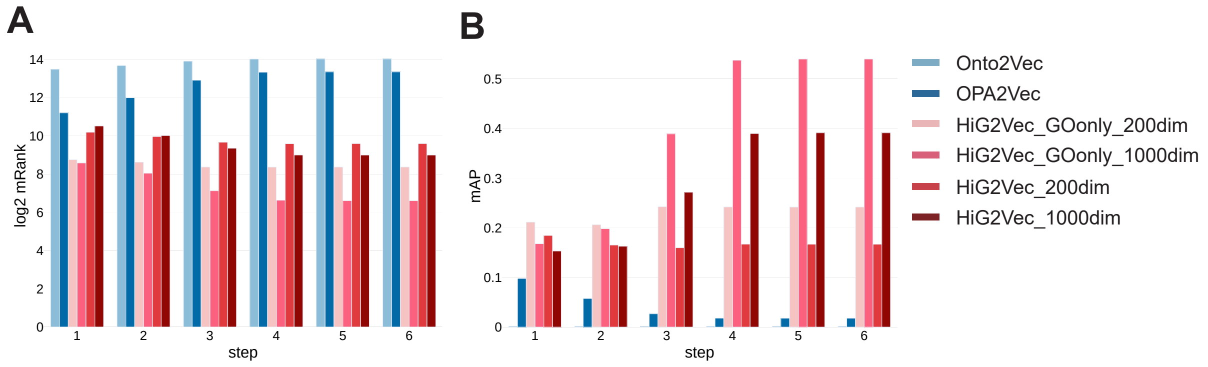


Figure S4: Hierarchy reconstruction for embeddings of yeast. (A) is log2 transformed mRank and (B) is mAP when reconstructing within n-step reachable nodes.

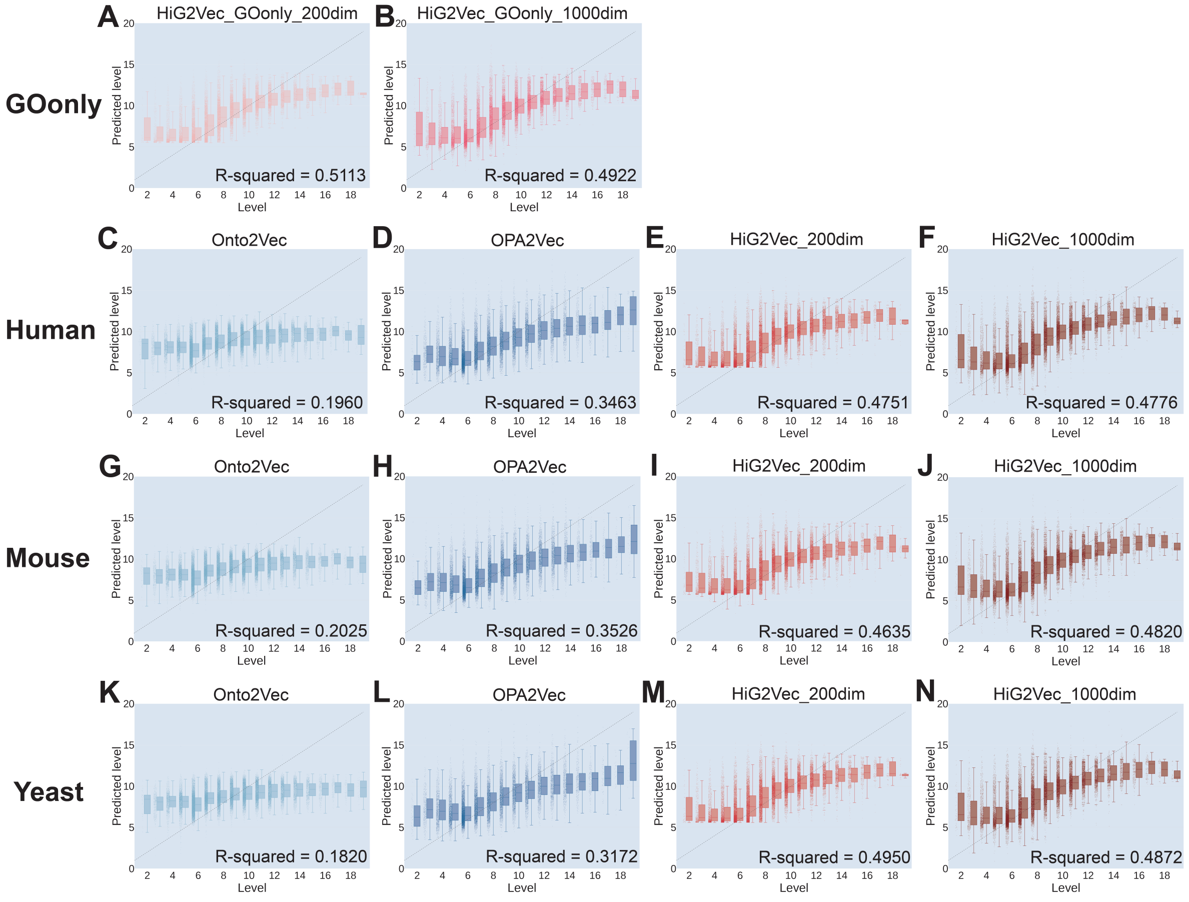


Figure S5: GO level prediction for every GO embeddings. (A,B) GOonly(using HiG2Vec), (C-F) Human, (G-J) Mouse, (K-N) Yeast

Table S5: Results of binary interaction prediction using the semantic similarity measures with BMA approach

| | Model | Domain | STRING_Human | STRING_Mouse | STRING_Yeast | HumanNet_XN |
|---|---|---|---|---|---|---|
| AUC | Resnik | BP | 0.8234 | **0.7329** | 0.8189 | 0.7799 |
| | | MF | 0.6379 | 0.6636 | 0.6531 | 0.6840 |
| | | CC | 0.7306 | 0.7114 | 0.8129 | 0.7194 |
| | Wang | BP | 0.8244 | 0.7242 | 0.8151 | 0.7779 |
| | | MF | 0.5871 | 0.6297 | 0.6714 | 0.6232 |
| | | CC | 0.7295 | 0.6952 | **0.8491** | 0.7122 |
| | GOGO | BP | **0.8483** | 0.7224 | 0.8252 | **0.7950** |
| | | MF | 0.4967 | 0.6331 | 0.7036 | 0.4984 |
| | | CC | 0.4979 | 0.7066 | 0.8439 | 0.4894 |

Table S6: STRING binary interaction prediction of the embedding methods

| | | | STRING_Human | | | HumanNet_XN | | |
|---|---|---|---|---|---|---|---|---|
| | | Dim | BMA | DIST | NN | BMA | DIST | NN |
| AUC | HiG2Vec | 10 | 0.8195 | 0.7670 | 0.8330 | 0.7799 | 0.7446 | 0.7858 |
| | | 20 | 0.8040 | 0.7869 | 0.8671 | 0.7683 | 0.7662 | 0.8114 |
| | | 50 | 0.8123 | 0.7887 | 0.9165 | 0.7740 | 0.7706 | 0.8432 |
| | | 100 | 0.8349 | 0.8102 | 0.9501 | 0.7940 | 0.7846 | 0.8746 |
| | | 200 | 0.8256 | 0.7843 | 0.9671 | 0.7836 | 0.7626 | 0.8872 |
| | | 500 | 0.7989 | 0.7270 | 0.9753 | 0.7607 | 0.7097 | 0.8917 |
| | | 1,000 | 0.7908 | 0.7027 | 0.9771 | 0.7538 | 0.6866 | 0.8944 |
| | OPA2Vec | 200 | 0.8267 | 0.7393 | **0.9773** | 0.8002 | 0.7563 | **0.9023** |
| | Onto2Vec | 200 | 0.6277 | 0.5218 | 0.7963 | 0.6300 | 0.5422 | 0.6243 |
| | Gene2Vec | 200 | - | 0.6138 | 0.8916 | - | 0.6476 | 0.7836 |

| | | | STRING_Mouse | | | STRING_Yeast | | |
|---|---|---|---|---|---|---|---|---|
| | | Dim | BMA | DIST | NN | BMA | DIST | NN |
| AUC | HiG2Vec | 10 | 0.7313 | 0.7048 | 0.7824 | 0.8548 | 0.7639 | 0.8223 |
| | | 20 | 0.7296 | 0.7068 | 0.8346 | 0.8528 | 0.7629 | 0.8817 |
| | | 50 | 0.6985 | 0.6499 | 0.8942 | 0.8630 | 0.7887 | 0.9397 |
| | | 100 | 0.7361 | 0.7118 | 0.9304 | 0.8690 | 0.8045 | 0.9637 |
| | | 200 | 0.7273 | 0.6922 | 0.9557 | 0.8632 | 0.7828 | 0.9677 |
| | | 500 | 0.6965 | 0.6287 | 0.9721 | 0.8482 | 0.7514 | 0.9725 |
| | | 1,000 | 0.6920 | 0.6178 | 0.9755 | 0.8389 | 0.7356 | 0.9734 |
| | OPA2Vec | 200 | 0.7322 | 0.7619 | **0.9758** | 0.8485 | 0.7901 | **0.9860** |
| | Onto2Vec | 200 | 0.5634 | 0.5159 | 0.7833 | 0.7109 | 0.5150 | 0.7362 |

Table S7: STRING interaction score prediction of the embedding methods

| | | STRING_Human | | STRING_Mouse | | STRING_Yeast | | HumanNet_XN | |
|---|---|---|---|---|---|---|---|---|---|
| | Dim | R-squared | RMSE | R-squared | RMSE | R-squared | RMSE | R-squared | RMSE |
| HiG2Vec | 10 | 0.1011 | 303.86 | 0.1219 | 304.92 | 0.0641 | 294.50 | 0.0748 | 0.99 |
| | 20 | 0.1526 | 295.02 | 0.1636 | 297.59 | 0.1563 | 279.63 | 0.1070 | 0.79 |
| | 50 | 0.2165 | 283.68 | 0.2571 | 280.47 | 0.2763 | 258.98 | 0.1592 | 0.94 |
| | 100 | 0.2880 | 270.44 | 0.3286 | 266.64 | 0.3653 | 242.53 | 0.2376 | 0.90 |
| | 200 | 0.3738 | 253.61 | 0.4531 | 240.63 | 0.4627 | 223.15 | 0.2955 | 0.86 |
| | 500 | 0.5810 | 207.44 | 0.6408 | 195.01 | 0.5624 | 201.39 | 0.3376 | 0.84 |
| | 1000 | **0.6532** | **188.75** | **0.6843** | **182.84** | 0.5862 | 195.83 | 0.3601 | 0.82 |
| OPA2Vec | 200 | 0.5165 | 222.85 | 0.5786 | 211.23 | **0.5863** | **195.79** | **0.3674** | **0.82** |
| Onto2Vec | 200 | 0.3339 | 261.80 | 0.4643 | 268.33 | 0.4315 | 228.57 | 0.1559 | 0.94 |
| Gene2Vec | 200 | 0.4879 | 229.44 | - | - | - | - | 0.2335 | 0.90 |

Table S8: STRING interaction type prediction of the embedding methods

| | | STRING_Human | | | STRING_Mouse | | | STRING_Yeast | | |
|---|---|---|---|---|---|---|---|---|---|---|
| | Dim | Acc | Macro F1 | Micro F1 | Acc | Macro F1 | Micro F1 | Acc | Macro F1 | Micro F1 |
| HiG2Vec | 10 | 0.3453 | 0.3053 | 0.7133 | 0.4019 | 0.3278 | 0.7625 | 0.5966 | 0.2693 | 0.7537 |
| | 20 | 0.4395 | 0.3272 | 0.7536 | 0.4404 | 0.3319 | 0.7890 | 0.6600 | 0.3197 | 0.8104 |
| | 50 | 0.5539 | 0.3566 | 0.8141 | 0.5885 | 0.3653 | 0.8464 | 0.7575 | 0.3734 | 0.8938 |
| | 100 | 0.6562 | 0.4433 | 0.8623 | 0.7117 | 0.4580 | 0.8921 | 0.8311 | 0.3931 | 0.9278 |
| | 200 | 0.7489 | 0.4797 | 0.9032 | 0.7906 | 0.4741 | 0.9244 | 0.8851 | 0.4056 | 0.9490 |
| | 500 | 0.8417 | 0.5168 | 0.9419 | 0.8820 | 0.5299 | 0.9594 | 0.9073 | 0.4116 | 0.9603 |
| | 1000 | **0.8710** | **0.5292** | **0.9524** | **0.9101** | **0.5440** | **0.9692** | 0.9110 | 0.4125 | 0.9617 |
| OPA2Vec | 200 | 0.8019 | 0.5009 | 0.9247 | 0.8440 | 0.4913 | 0.9462 | **0.9224** | **0.4163** | **0.9682** |
| Onto2Vec | 200 | 0.6200 | 0.3732 | 0.8497 | 0.7090 | 0.4573 | 0.8913 | 0.8580 | 0.4005 | 0.9389 |
| Gene2Vec | 200 | 0.7230 | 0.3942 | 0.8968 | - | - | - | - | - | - |
